# The genomic and transcriptomic foundations of viviparous seed development in mangroves

**DOI:** 10.1101/2020.10.19.346163

**Authors:** Hongmei Qiao, Xiaoxuan Zhou, Wenyue Su, Xing Zhao, Pengfei Jin, Shanshan He, Wei Hu, Meiping Fu, Dingtian Yu, Saiqi Hao, Yuan-Ye Zhang, Wenqing Wang, Congting Ye, Qingshun Quinn Li, Yingjia Shen

## Abstract

Vivipary in plants refers to a specific seed development and reproductive strategy where seeds minimize the dormancy stage and germinate while still attached to their maternal plants. It is one of the most unique adaptive genetic features used by many mangrove species where elongated hypocotyls aid in quick root emergence to anchor the seedling in coastal intertidal wetlands. The genetic mechanisms behind mangrove vivipary, however, remain elusive. Using comparative genomic and transcriptomic technologies to investigate viviparous mangroves and their close inland relatives, we found that a full array of gene expression profiles were altered, including key plant hormone metabolic pathways, high expression of embryonic signature genes, and reduced production of proanthocyanidins and storage proteins. Along with these changes, a major gene regulating seed dormancy, *Delay of Germination-1* (*DOG1*), is entirely missing or defunct within the entire linage of the four genera with true viviparous characteristics. These results suggest a systemic level change is required to warrant the genetic program of mangrove vivipary. Understanding of the molecular processes of vivipary could benefit the design of pregerminated propagules for forestation in harsh environments or prevent precocious germination of grain crops pre- and post-harvest.

## Introduction

Seed development, dormancy and germination are necessary reproductive processes shared by most plant species. In vivipary plants, seed development skips dormancy and proceeds directly into the germination phase where embryos, through an innate genetic program, continue to grow and germinate into extensive hypocotyls before dispersal (Elmqvist and Cox 1996). This specific reproductive model is noteworthily found in four genera of the Rhizophoraceae family of mangroves, *Kandelia, Rhizophora, Bruguiera*, and *Ceriops*, whose habitats are located in the tropical and subtropical intertidal zones (Tomlinson 2016). Studies show that the Rhizophoraceae family originated at about 54 million years ago (Xu et al. 2017) and developed many features adaptive to intertidal zone at the genomic level (Xu et al. 2017; He et al. 2020b). However, the genetic and molecular mechanisms behind the vivipary phenomenon are still unclear.

Phytohormones Abscisic Acid (ABA) and Gibberellic Acid (GA) are recognized as master regulators that play antagonistic roles in seed dormancy and germination across plant species (Shu et al. 2016; Liu and Hou 2018). Genes related to ABA biosynthesis (*ZEP*/*ABA1, NCED, SDR*/*ABA2, AAO, ABA3*) (Chen et al. 2020), inactivation/degradation (*CYP707A*) (Kushiro et al. 2004) and response (*ABI3, ABI4, ABI5*) (Brocard-Gifford et al. 2003) determine the activities of ABA. Previous reports have shown that both embryonic and maternal ABA contents in mangrove plants are lower than those in non-mangrove plants (Farnsworth and Farrant 1998). Maternal ABA content also exceeds embryonic tissues in Rhizophoraceae, suggesting a functional trade-off between the maintenance of embryonic development and tolerance to salinity or other stressors (Farnsworth and Farrant 1998). External applications of ABA in high concentrations could also repress vivipary (Hong et al. 2018). In the GA metabolism pathway, a series of genes, *KS, KO, KAO, GA20OX, GA3OX* and *GA2OX* control the level of GA (He et al. 2020a). Proanthocyanidins (PAs) are a collection of secondary metabolites also known to play roles in inhibiting seed germination by reinforcing the strength of the seed coat with increased levels of pigmentation and reduced water permeability (Debeaujon et al. 2000; MacGregor et al. 2014). Seeds of PA synthesis mutants were less dormant (Haughn and Chaudhury 2005).However, studies on the effects of ABA, GA and PA on vivipary are very limited.

In addition to plant hormones, genes that participate in seed dormancy and germination may also affect vivipary. It is widely accepted that the *DOG1* gene in *Arabidopsis* encodes a key promoter/inhibitor of seed dormancy/germination (Carrillo-Barral et al. 2020). *DOG1* and its four family members, called *DOG1 like1-4*, were also identified in many other species such as wheat, barley, rice, maize and *Brachypodium* (Ashikawa et al. 2013) (Fig. S1). Studies indicated that *DOG1* controls not only the dormancy of orthodox seeds but also the germination time of recalcitrant seeds through interactions with ABA and GA (Chiang et al. 2011; Nakabayashi et al. 2012; Graeber et al. 2014; Nonogaki 2019). In *Arabidopsis, DOG1* acts in parallel with ABA by affecting *PP2C*s, thus enhancing ABA sensitivity (Née et al. 2017), while the *dog1* mutant showed non-dormancy and increased GA sensitivity (Bentsink et al. 2006; Nakabayashi et al. 2012). The effects of *DOG1* on seed germination are also demonstrated through its regulation of *GA20OX*, a GA biosynthesis gene, and genes encoding downstream cell-wall remodeling proteins all directly related to seed germination in a temperature dependent manner (Graeber et al. 2014). However, the function of *DOG1* in woody plants, including its role in mangrove vivipary, is unknown.

Adequate nutrients stored in seeds are essential for germination. To this end, seed storage proteins, maturation proteins and late embryogenesis abundant proteins are massively accumulated during seed development, especially at the late stage of maturation and act as the main source of nitrogen, carbon, and sulfur at germination(Wei et al. 2020). However, lack of nutrient storage in vivipary mangrove embryos could influence the development of seeds, promoting germination using nutrients from maternal tissues.

To exploit the genetic mechanism of vivipary, we sequenced the genomes of three vivipary species [*Kandelia obovata* Sheue, Liu & Yong (Sheue et al. 2003), *Bruguiera gymnorhiza* (L.) Savigny and *Ceriops tagal* (Pers.) C. B. Rob.] and one of its non-viviparous relatives [*Carallia brachiata* (Lour.) Merr.], an inland species of the Rhizophoraceae that produces orthodox seeds with dormancy (Schwarzbach and Ricklefs 2000; Shi et al. 2002). Moreover, we also sequenced the transcriptomes of the reproductive organs in *K. obovata* and *C. brachiata* through a series of stages during seed development. After comparative genomic and transcriptomic analyses and confirmation, we demonstrated that the contraction of key gene families and the subsequent hormonal changes are directly related to the distinct phenotype of vivipary in mangroves.

## Results

### Assembly and annotation of *K. obovata* and *C. brachiata* genomes

The genome of *K. obovata*, estimated at ∼224 Mb in size, was first assembled using PacBio, 10X Genomics, and Illumina paired-end sequencing data. The resulting genome assembly consists of 166,469 contigs (with a scaffold N50 >750kb; total assembled size 209.87Mb; Table 1). The Hi-C technology was further used to order and orient 353 contigs into 18 pseudo-chromosomes (Fig. 1, Fig. S2 and Supplementary Table S1). The *C. brachiata* genome, ∼364 Mb in size, was done by PacBio and Illumina sequencing, and consisted of 729 scaffolds (with a scaffold N50 of > 1.53 Mb; assembly size 341.48 Mb; Table 1). The accuracy of the genomes was evaluated by mapping and coverage rates at 99.39% and 95.62% for *K. obovata*; 97.15% and 98.08% for *C. brachiata* respectively (Supplementary Table S2). The reference genome of *K. obovata* has 23,683 protein-coding genes, which is significantly less than that in *C. brachiata* (32,279) and other woody plants (Supplementary Table S3). Two additional draft genomes of viviparous mangroves, *B. gymnorhiza* and *C. tagal*, were also sequenced, assembled and polished using Oxford Nanopore and Illumina reads (Table 1).

**Table 1.** Statistics of genomes assembled

| Level | Statistics | *K. obovata* | *C. brachiata* | *C. tagal* | *B. gymnorhiza* |
| --- | --- | --- | --- | --- | --- |
| Contig | Number | 166,469 | 290,222 | 1,514 | 3,331 |
|  | Longest Length (bp) | 4,026,005 | 505,877 | 9,673,137 | 6,128,382 |
|  | Total size (bp) | 209,874,737 | 338,878,113 | 259,034,946 | 266,782,858 |
|  | N50 length (bp) | 750,717 | 34,575 | 2,934,242 | 806,366 |
| Scaffold | Number | 165,418 | 279,914 | - | - |
|  | Longest Length (bp) | 15,414,410 | 6,734,800 | - | - |
|  | Total size (bp) | 210,614,605 | 341,477,579 | - | - |
|  | N50 length (bp) | 5,432,096 | 1,078,353 | - | - |
| Chromosome | Number | 18 | - | - | - |
|  | Longest Length (bp) | 14,017,977 | - | - | - |
|  | Total size (bp) | 172,840,403 | - | - | - |
|  | N50 length (bp) | 10,005,181 | - | - | - |

**Fig. 1.**
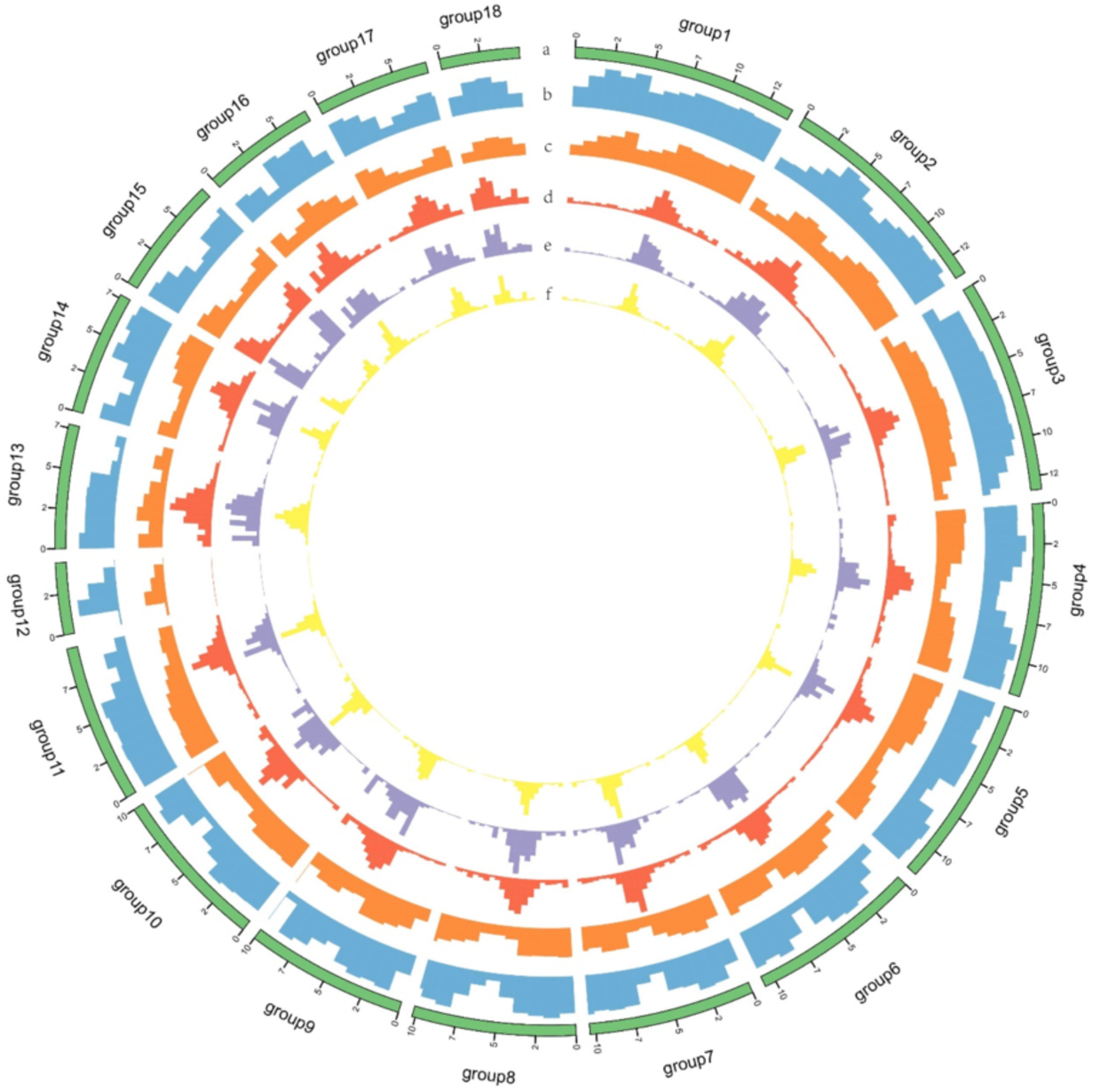
Circular diagram depicting the characteristics of the *K. obovata* genome. a, The pseudochromosomes. b, Gene density in 1Mb sliding windows. c, GC-content in 1Mb sliding windows. d, repeat density in 500kb sliding windows. e, *copia* density in 500kb sliding windows. f, *gypsy* density in 500kb sliding windows.

### Phytohormone ABA dynamics accompany vivipary development

It was previously shown that phytohormones ABA and GA are important for vivipary development (Xu et al. 2017), we thus examined the expression profiles of hormone related genes during crucial stages of viviparous embryo development and seed germination. In *K. obovata* (Fig. 2A), an ovule (termed Stage 1 or S1) is fertilized, then a liner-shaped embryo is formed (S2). Following cotyledon filling of the embryo sac (S3), the differentiated axis continues to elongate out of the seed coat (S4). The transition from S3 to S4 marks viviparous germination where the radicle breaks the seed coat.

**Fig. 2.**
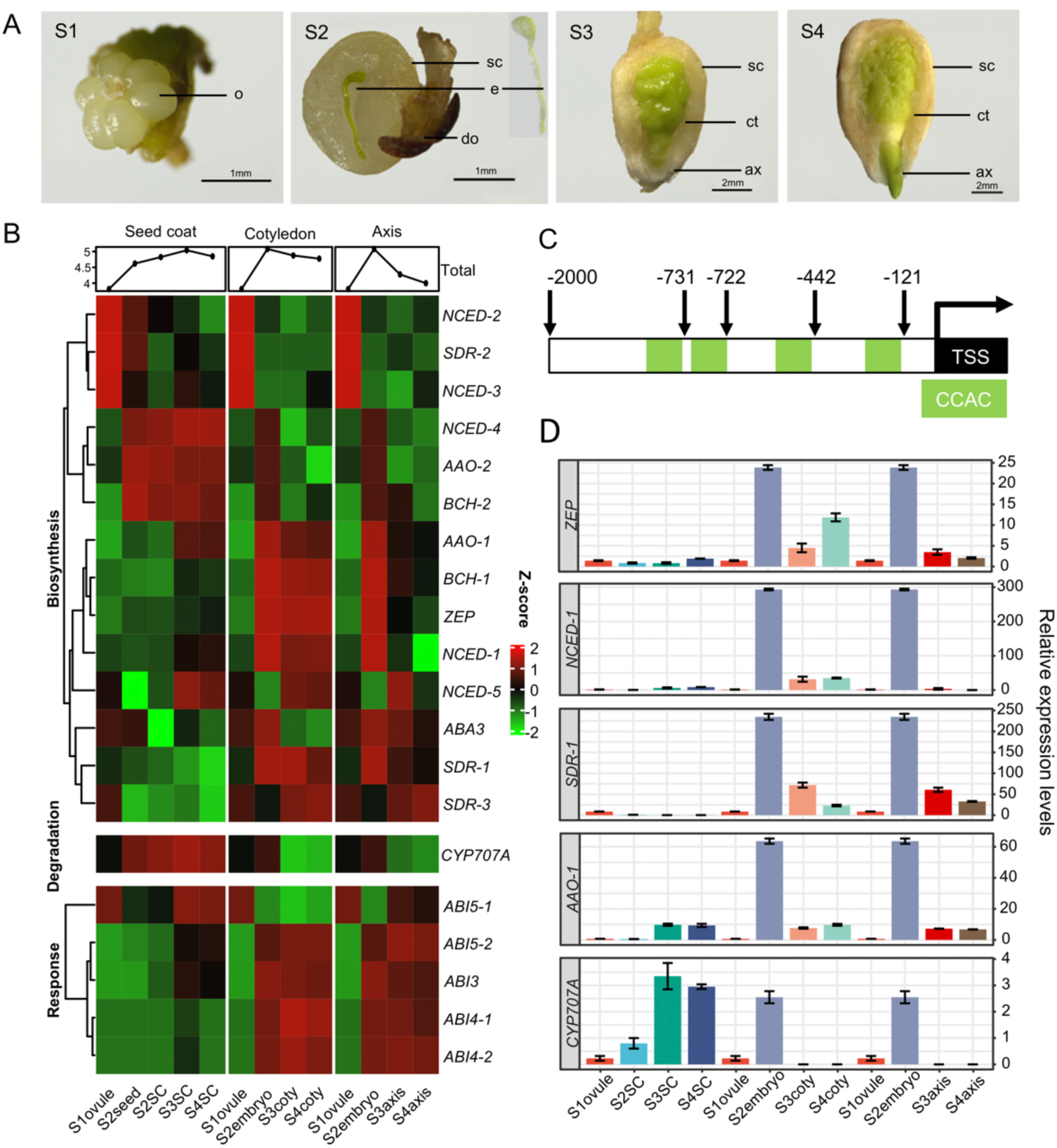
Dynamic of ABA during viviparous developmental stages. **A**. Morphological observation during viviparous process. S1, Stage1 unfertilized ovules. S2, fertilized seed with liner-shaped embryo (inset). S3, developing embryo in a seed sac. S4, axis protrude out of testa (seed coat). ax, axis; ct, coty, cotyledon; do, dead ovule; e, embryo; o, ovule; sc, seed coat. **B**. RNA-seq expression profiles of ABA metabolic and responsive genes. These 20 genes were divided into three clusters according to their functions. The Z-score scale shown on the right. **C**. Potential *ABI4* binding motifs (green) in *CYP707A* promoter; numbers mark CCAC motif locations in upstream of the transcription start site (TSS). **D**. RT-qPCR results of key genes in ABA metabolic. Relative expression levels are marked on the right; error bar represents standard deviation. For comparison purposes, S1ovule and S2embryo are shown more than once.

To probe into gene expression in different anatomical parts, viviparous seeds in S2 to S4 were dissected and separated into seed coat and embryo (further into cotyledon and axis in S3 and S4) for further transcriptome sequencing and analyses. Genes related to ABA biosynthesis (*ZEP*/*ABA1, NCED, SDR*/*ABA2, AAO, ABA3*) (Debeaujon et al. 2000), response (*ABI3, ABI4, ABI5*) (MacGregor et al. 2014), and inactivation/degradation (*CYP707A*) (Haughn and Chaudhury 2005), were extracted and displayed in Fig. 2B. A clear spatial-temporal pattern, along with the expression trend of these genes from embryo development to germination, were shown in three groups.

In the biosynthesis related group, seed coat and cotyledon were the main tissues that generated ABA. Three genes, namely *NCED-2, NCED-3* and *SDR-2*, were exclusively expressed in the S1 ovule, while *NCED-4, AAO-2* and *BCH-2* were highly expressed in seed coats. *AAO-1, BCH-1, ZEP, NCED-1, ABA3* and *SDR-1* were preferentially expressed in S2 but drastically decreased in S3 and S4 embryos. However, the availability of the ABA pool is determined by levels of biosynthesis and inactivation (modification and degradation). The degradation gene *CYP707A* was highly expressed in the S3 and S4 seed coats, thus reducing ABA concentration. In the ABA responsive genes group, except *ABI5-1*, transcription factors *ABI5-2, ABI3, ABI4-1*, and *ABI4-2*, mainly located in the cotyledon and axis, were expressed concordantly with ABA biosynthesis genes, suggesting their synergistic roles in this process.

The spatial specific expression of *CYP707A* in the seed coat may be a consequence of low *ABI4* expression in the same tissue. It has been demonstrated in *Arabidopsis* that *ABI4* suppresses the promotor activity of *CYP707A* by targeting its CCAC motifs (Shu et al. 2013). Interestingly, similar motifs have been found in the *CYP707A* promoter region in *K. obovata* (Fig. 2C). Thus, it is very likely that reduced expression of *ABI4* releases the suppression and increases *CYP707A* levels in the seed coat.

Expression of *ZEP, NCED-1, SDR-1, AAO-1*, and *CYP707A* were validated by using RT-qPCR (Fig. 2D). Consistent with results from RNA-seq, all biosynthesis genes climaxed in the S2 embryo and then dramatically decreased in later stages. The expression of *CYP707A* in seed coats reached a climax in S3, then reduced in S4. These results support the notion of ABA levels dramatically decreasing during the transition to vivipary.

### GA profile promotes vivipary

The comparison of transcriptomes between *K. obovata* and *C. brachiata* showed that GA biosynthetic genes (*KS, KO, KAO, GA20OX* and *GA3OX*) and positive transduction genes (*GID1* and *PIF4*) are up-regulated in *K. obovata*, while the negative transduction gene *DELLA* is down-regulated (Fig. 3A). The high expression of biosynthetic genes suggests that the *K. obovata* embryo produces higher contents of GA which facilitate viviparous germination. Among the signal transduction genes, *PIF4* could promote hypocotyl elongation, which may also promote the initiation of vivipary.

**Fig. 3.**
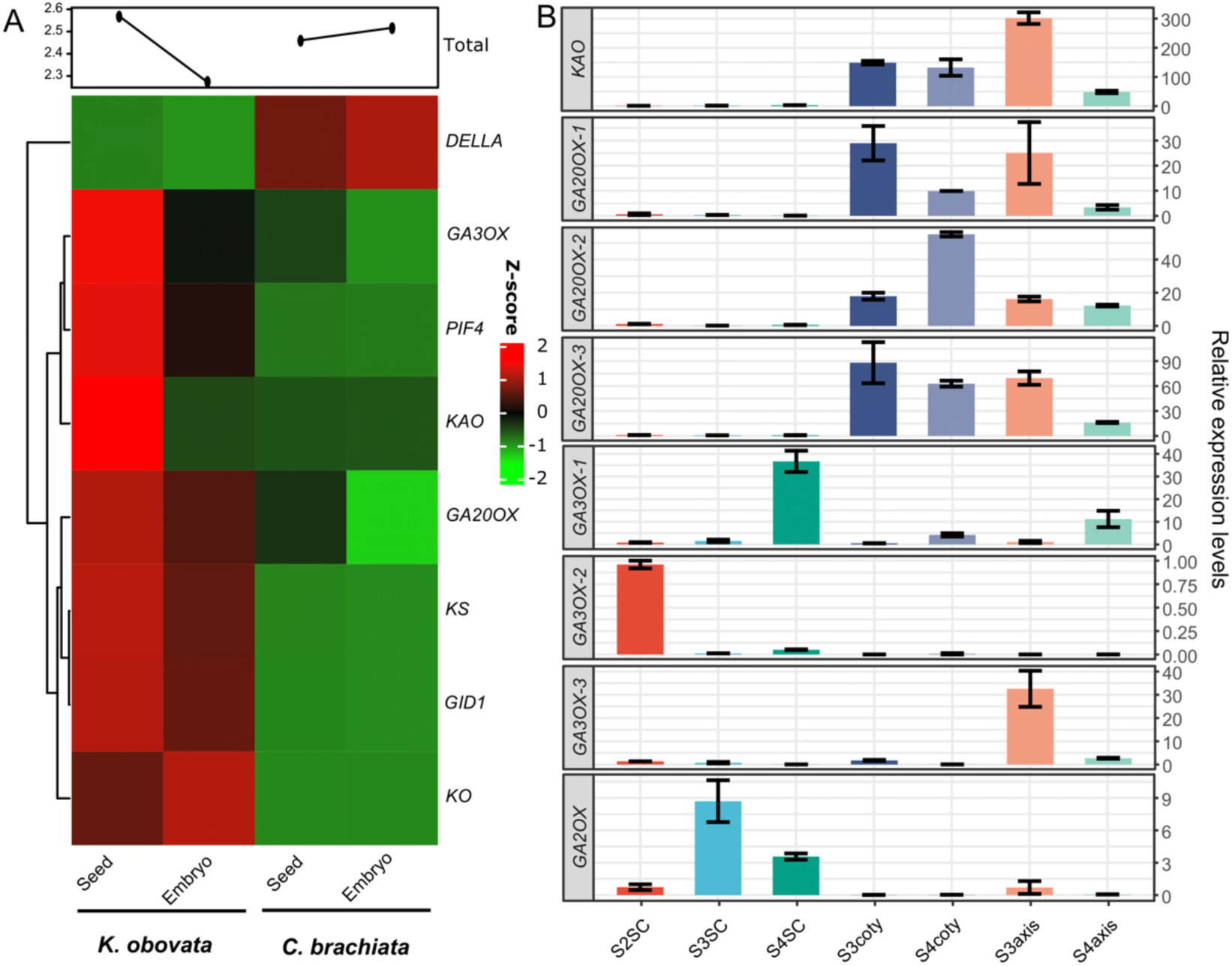
Expression profile of key genes in GA synthetic and signaling pathways in *C. brachiata* and *K. obovata*. **A**. Comparative expression levels of GA related genes in seed/embryo between two species by Z-score. **B**, RT-qPCR results of key genes in GA metabolic in viviparous tissues of *K. obovata*. Relative expression levels are marked on the right; error bar represents standard deviation. Coty, cotyledon; SC, seed coat. *GA20OX*, GA 20-oxidase; *GA3OX*, GA 3-oxidase; *GID1*, GA insensitive dwarf1; *KS, ent*-kaurene synthase; *KO, ent*-kaurene oxidase; *KAO, ent*-kaurenoic acid oxidase; *PIF4*, phytochrome interacting factor 4.

To probe into the role of GA in vivipary, expressions of GA metabolic genes were confirmed by RT-qPCR analyses (Fig. 3B). The GA biosynthesis genes *KAO, GA20OX-1* and *GA20OX-3* showed a consistent higher expression in S3 embryonic tissues (cotyledon and axis), especially *GA3OX-3*, which presented extremely high expression in the S3 axis. *GA20OX-2* and *GA3OX-1* were higher in S4 embryo and seed coat, while *GA3OX-2* was at a relatively lower level. When it comes to catabolism, *GA2OX* was preferentially expressed in the S3 and S4 seed coats rather than in the embryonic tissues. In general, up regulation of GA biosynthesis genes promotes the growth of germinating embryos, particularly in the axis.

To promote germination, GA also exhibits its effect through cell wall modification and expansion. Among the 102 cell-wall remolding proteins including amylase, chitinase, and expansion that were induced by GA (Nonogaki et al. 2007), a majority of the genes were expressed higher in S3SC (Fig. S3) where fast cell expansion is needed.

These results imply that high levels of GA were maintained to promote continual embryo growth and overcome dormancy.

### Proanthocyanidin reduction facilitates vivipary

PAs in seed coats generally prevent germination under unfavorable conditions. To investigate the role of PAs in vivipary, we extracted the expression profile of the genes encoding PA biosynthetic enzymes (Tian et al. 2007) in *K. obovata* and *C. brachiata*. The majority of the enzymes in PA synthesis were expressed higher in the embryos and seeds of *K. obovata* than those in *C. brachiata*. However, two specific genes encoding key enzymes near the end of pathway, *ANR* and *LAC15*, have higher expressions in *C. brachiata* (Fig. 4A and B). It was reported that the yellow seeds of *ANR*/*BAN* mutants in *Arabidopsis* have less seed coat-imposed dormancy and require less GA to germinate (Debeaujon et al. 2003). Therefore, the lower expression of *ANR* in *K. obovata* might be the main regulatory gene that reduces seed coat-imposed dormancy. *LAC15* colocalized with the flavonoid end products, proanthocyanidins and flavanols, when it was expressed in developing testa, thus making the seed coat brown (Pourcel et al. 2005), as seen in *C. brachiata* (Fig. S4).

**Fig. 4.**
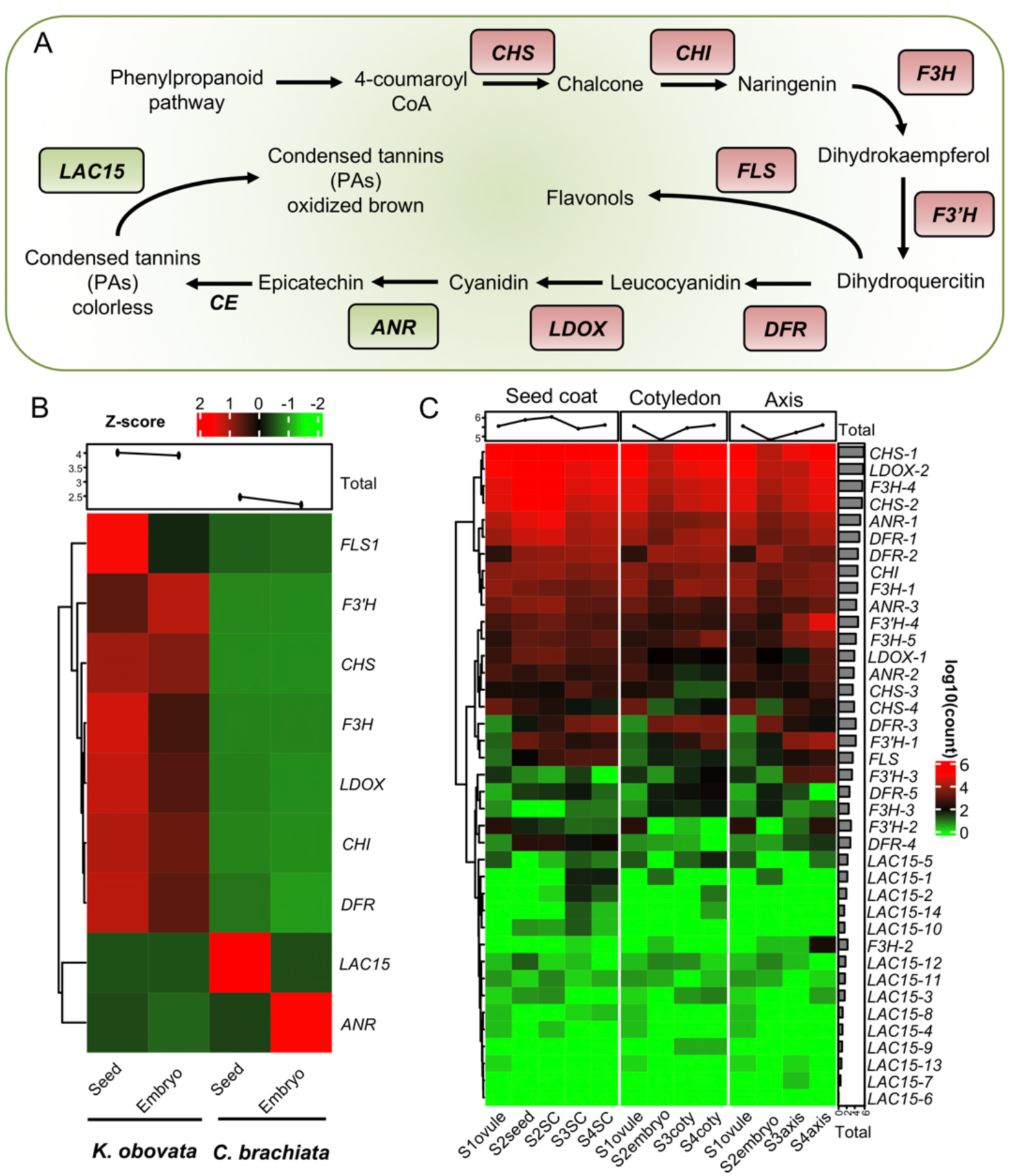
Proanthocyanidin pathway genes expression profiles in *K. obovata* and *C. brachiata*. **A**, Synthetic pathways of proanthocyanidins, enzymes catalyzing each reaction are listed next to the arrows, red and green highlight genes higher expressed in either *K. obovata* or *C. brachiata*. **B**, Expression profiles of synthetic genes of proanthocyanidin pathway between *K. obovata* and *C. brachiata*. Z-score scale is shown on the top. **C**, Expression profiles of proanthocyanidins biosynthetic genes in viviparous tissues. Colors represent log10 of the reads in RNA-seq. Coty, cotyledon; SC, seed coat. *FLS*, flavanol synthase; *F3’H*, flavonoid 3’ hydroxylase; *CHS*, chalcone synthase; *CHI*, chalcone isomerase; *F3H*, (2S)-flavanone 3-hydroxylase; *DFR*, dihydroflavonol 4-reductase; *LDOX*, leucoanthocyanidin dioxygenase; *ANR*, anthocyanidin reductase; *CE*, condensing enzyme; *LAC*, Laccases.

During vivipary, PAs were highly expressed in seed coats when compared to embryonic tissues, reaching a peak in S2 and then decreasing. The S2embryo possessed the lowest levels of these biosynthesis gene expression, which increased in S3 and S4 (Fig. 4C). Another noticeable fact was that the whole family of *LAC15* genes were almost unexpressed in all stages and tissues. Given the key function of *LAC15*, its silencing may cause white seed coat of *K. obovata* (Fig. 2A). These results suggest that such a reduction of PA production in the seed coat may directly loosen the seed coat in S3 and S4, thus facilitating axis protrusion and hence vivipary.

### Genetic bases of dormancy loss

To investigate the genetic mechanism of vivipary, 729 genes or gene families that were reported to participate in seed development, dormancy and germination in *Arabidopsis* were listed as candidates for further interrogation (Supplementary Table S4). Through detailed gene-level homology comparisons, homologs of most genes were identified in the genome of *K. obovata*. However, it was found that four gene families were exclusively lost in *K. obovata* and *R. apiculata* at both genomic and transcriptomic levels. Among them, *DOG1* is the most conspicuous for its decisive role in determining seed dormancy (more detail in Methods: Identification of missing genes for vivipary) and the germination of seeds, while also being suggested as a repressor of GA biosynthesis (Skubacz and Daszkowska-Golec 2017). Since *DOG1* is missing in viviparous mangroves, we therefore postulate that the increase of bioactive GA from non-vivipary to vivipary stages promotes seed germination.

To further interrogate the genetic foundation leading to vivipary, detailed analysis of *DOG1* in Rhizophoraceae species was performed. The initial search for the *DOG1* homologue in the *K. obovata* genome was not fruitful, but a homologue was clearly shown in the genome of *C. brachiata*. Comparison based on the genomic regions of six genes upstream and five genes downstream of the *DOG1* locus in *C. brachiata* revealed that the order of these genes are conserved between *C. brachiata* and *K. obovata*, except for a deletion of the *DOG1/DOG1-like* genic region in the latter (Fig. 5B). To expand the search of *DOG1* in all four viviparous genera of Rhizophoraceae, we downloaded the genome of *R. apiculata* (Xu et al. 2017) in addition to genomes of three viviparous species for comparison. As shown in Fig. 5B, among these six species (4 viviparous, and 2 non-viviparous species: *C. brachiata* and *Manihot esculenta* Crantz), the functional *DOG1* gene only exists in two terrestrial trees but is completely lost in *C. tagal, K. obovata*, and *R. apiculata*, while *B. gymnorrhiza* retains an altered non-functional copy.

**Fig. 5.**
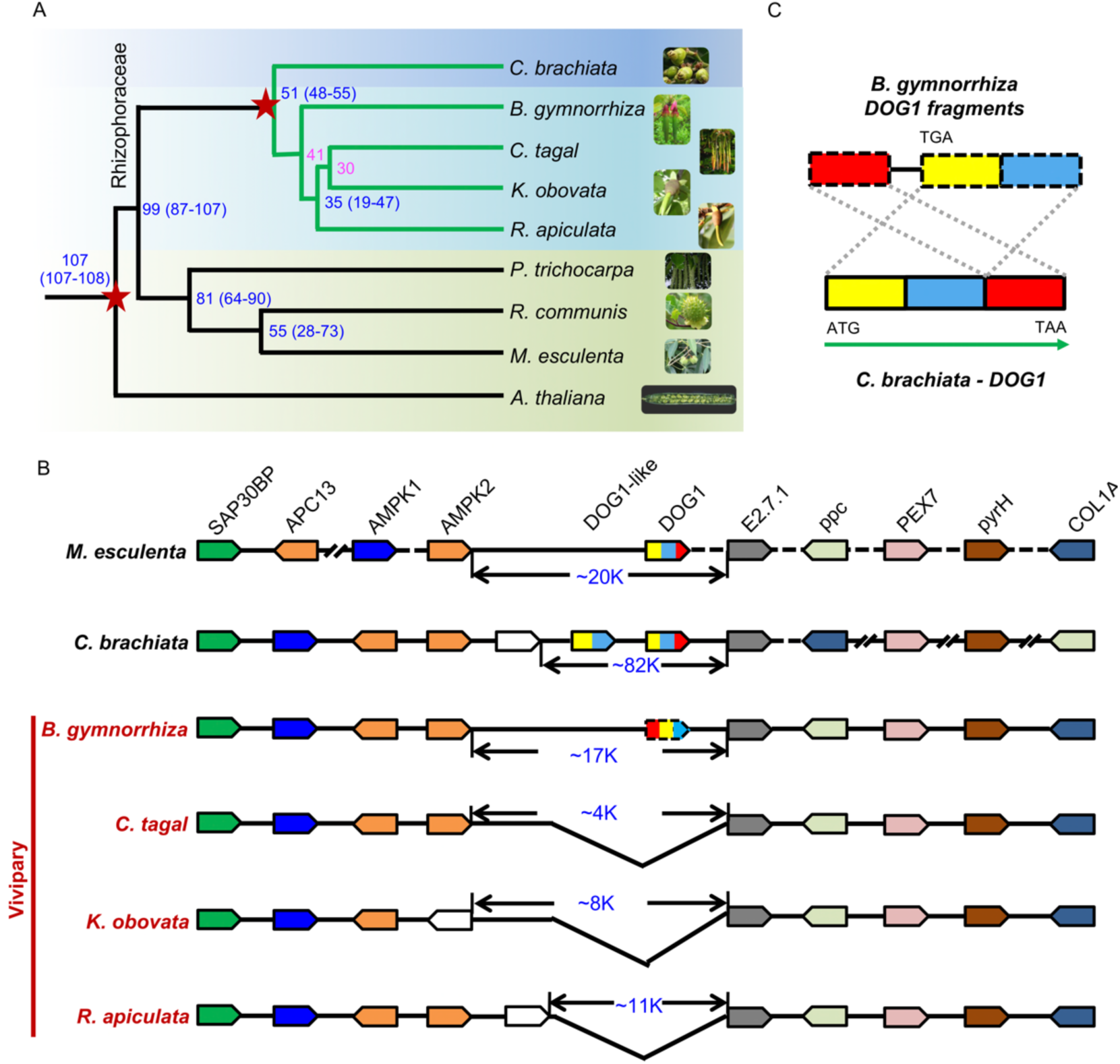
Phylogenetic analysis and the synteny of *DOG1* genomic regions. **A**, Phylogenic analysis of vivipary mangrove and related plants. Numbers are divergence times and confidence intervals (in parentheses, million years ago). The pink divergence times and calibration points (red stars) were cited from Xu et al. 2017. **B**, Synteny of the genomic regions around *DOG1*. Colored polygons indicate genes and arrowheads show transcriptional orientations. Double slanted black lines, genes on either side are not directly linked. Black dashed lines mean some other genes there. **C**, Alignment of *DOG1* between *B. gymnorrhiza* and *C. brachiata*. Each color represents a domain. Boxes with solid lines mean coding sequence of *DOG1;* boxes with dash lines indicate incomplete CDS lacking a start codon.

The alignment of *DOG1* sequences between *B. gymnorrhiza* and *C. brachiata* indicate that three domains were reversed in *B. gymnorrhiza*, a start codon being only present in the second domain, but not the first (Fig. 5C and Fig. S5). There was also an insertion fragment of ∼600 bp between the first two domains (Fig. 5C). All of these changes destroyed the gene structure and made it a pseudogene in *B. gymnorrhiza*.

Further analyses show that genes upstream and downstream of the *DOG1* locus were relatively conserved, especially the upstream genes located on the same chromosome with conserved synteny among *C. brachiata, B. gymnorrhiza, C. tagal, K. obovata* and *R. apiculata*, but not in *M. esculenta*. The five genes downstream of *DOG1* were located on the same chromosome in *M. esculenta, B. gymnorrhiza, C. tagal, K. obovata* and *R. apiculata*, but not in *C. brachiata*. The most important finding, however, was that the *DOG1* gene was nonfunctional or completely lost between *AMPK* and *E2*.*7*.*1* in all four viviparous species (Fig. 5B). The sizes of the deleted sequences between these two genes ranged from 4K to 17K.

To eliminate the possibility that the *DOG1* regions were not assembled successfully in mangroves, raw PacBio/Nanopore and Illumina reads used for genome assembly of viviparous mangroves were mapped to the *DOG1* region of *C. brachiata* and yet no alignment was found (Fig. S6). In contrast, there is a functional copy of *DOG1* in each of the two non-viviparous species, *M. esculenta* and *C. brachiata*. There is another *DOG1-like* gene which only contains the first two domains of *DOG1* in *C. brachiata* but is missing in all other species (Fig. 5B, Fig. S5A and B). Tissue specific expression analysis showed that the *DOG1* is only expressed in the fruit/seed/embryo (reproductive organs) of *C. brachiata* but missing in *K. obovata* (Fig. S5D).

To verify the function of *C. brachiata DOG1*, we transferred the *DOG1* of *C. brachiata* (*CbDOG1*) and *A. thaliana* (*AtDOG1*, as a positive control) into the *dog1-5* mutant background [which showed the phenotype of deep dormancy; (Fedak et al. 2016)] and tested seed germination rate in the transgenic Arabidopsis. The results showed that the transgenic line *dog1-5+CbDOG1* and *dog1-5+AtDOG1* had comparable seed germination rates with the wide type *Col-0*, but had a significantly higher germination rate than *dog1-5* (Fig. S7). Thus, we conclude that *CbDOG1* and *AtDOG1* are similar in function, which is indicative of an evolutionary conservation, and that the missing *DOG1* gene in the Rhizophoraceae family of mangroves could contribute to the loss of dormancy.

### Lack of seed storage proteins in viviparous mangroves

Nutrient storage functions in preparation for future germination of orthodox seeds. However, due to the constant supply of nutrients from the maternal tree, such a storage may not be necessary in viviparous seeds. To resolve this, we investigated the seed protein genes in the genomes of seven species, including two viviparous (*K. obovata* and *R. apiculata*) and 5 non-viviparous species (*C. brachiata, A. thaliana, M. esculenta, P. trichocarpa*, and *R. communis*). The results showed that most protein genes were only missing in the genomes of two viviparous species (Supplementary Table S5). Moreover, they were specifically and highly expressed in the reproductive organs of *C. brachiata*. Among many other functions, seed storage proteins (orthogroup OG0001910) serve as sources of carbon and nitrogen for seed germination (Wei et al. 2020). Oleosin family proteins (orthogroups: OG0002151 and OG0007120) are another nutrient in seeds (Lu et al. 2018) while seed maturation proteins (orthogroup: OG0014483 and OG0015329) are involved in secondary dormancy (Kushwaha et al. 2012). Late embryogenesis abundant proteins (LEA) (orthogroup: OG0009552, OG0012500 and OG0012688) play an important role in the desiccation tolerance of plant seeds (Pizarro et al. 2019). Thus, there are clear differences within these sets of proteins in terms of their coding genes and expressions between viviparous mangroves and orthodox seeds. Such a genomic level difference may lead to the germination of viviparous seeds while attached to mother trees.

## Discussion

Vivipary is a specific seed development strategy and a notable feature of many mangrove species. However, the molecular mechanism of vivipary was never revealed, partly due to limited genome information and difficulties in dissecting crucial development stages. Previous studies were mostly focused on physio-biochemical level characterizations. In our study, the hormonal dynamics at these two crucial stages of transition, from embryo development to germination, showed that elevated levels of GA/ABA and loosened seed coats promoted germination and axis growth and were a result of reduced PA. In *Arabidopsis*, it was shown that GA biosynthesis genes were highly expressed after imbibition in embryos especially in radicles (Ogawa et al. 2003; Penfield 2017), while ABA was produced in endosperms and embryos during seed development and maturation (Rodríguez-Gacio et al. 2009). Their biosynthesis activities were separated by seed dehydration and dormancy. However, such a clear division is no longer observed in *K. obovata*, where both ABA and GA syntheses happen side-by-side in the same tissues of cotyledon and axis (Figs. 2 and 3). In contrast, GA synthesis genes are expressed at low levels in *C. brachiata* (Fig. 2A), ensuring the separation of GA and ABA biosynthesis. These results imply that there may not be a clear boundary between seed development and germination in viviparous mangroves; the high GA/ABA ratio is set up before germination occurs.

Upon systemic searches of the genetic foundation of vivipary, we identified the loss of a key gene controlling seed dormancy and germination, *DOG1;* which may have undergone two continuous rounds of mutations in the lineage of viviparous mangrove species. The first event could have happened to *DOG1* when a fragment was inserted in to the gene, destroying the reading frame in a basal mangrove species, *B. gymnorrhiza*. The non-functional *DOG1* lost its ability of encoding a protein, which not only produced non-dormancy seeds but also opened the door for germination. Such an evolved trait might have made viviparous mangroves capable of surviving in intertidal habitats, possessing great advantages that were favorably selected by nature. The second event that happened to the *DOG1* could be that the remaining body of this gene was completely removed from the genomes in a new genus, *Rhizophora*. Newer viviparous mangroves, such as two newly differentiated genus *Kandelia* and *Ceriops*, all lost *DOG1* in their genomes. The evolutionary route of this gene was also proven by a phylogenetic study of a mangrove’s genome (Xu et al. 2017) (Fig. 5A).

Storing enough nutrients during seed development is essential for seed germination after separation from maternal trees. Our study shows that genes related to energy storage were lost in the viviparous genomes. However, the causal reason is unclear. We speculate that, due to a lack of nutrient storage, these viviparous mangrove seeds germinate while attached to mother trees. An alternative theory would be that adapting to viviparous situations, the energy storage related genes were not selected. However, such a massive loss of genes would not have happened without dreadful selection pressures. Thus, further study is necessary to resolve this issue.

In summary, a system level change of in mangrove lineage represented by *K. obovata* leading to the adaptive features of vivipary. As depicted in Figure 6, these changes include the deletion of the *DOG1* gene, the alteration of ABA/GA dynamics, the softening of the seed coat by PA content reduction, and the loss of energy storages in seeds. Decrease in ABA during seed maturation may directly cause a loss in dormancy. Increases in the ratio of GA/ABA from non-vivipary to vivipary and declining PA content in surrounding tissues and subsequently permits germination to occur. The lack of nutrient storage might prompt the selection of propagules that could grow embryos with lengthened hypocotyls in order to increase energy reservations when dispersed. While the connections among these theories are not solid at this time, our study provides a comprehensive outline for future research on vivipary. Studying the molecular mechanisms of vivipary in mangroves provides a unique opportunity to better understand seed dormancy, prevent pre-harvest germination in crops, and understand the adaptation of mangroves to the coastal environment.

**Fig. 6.**
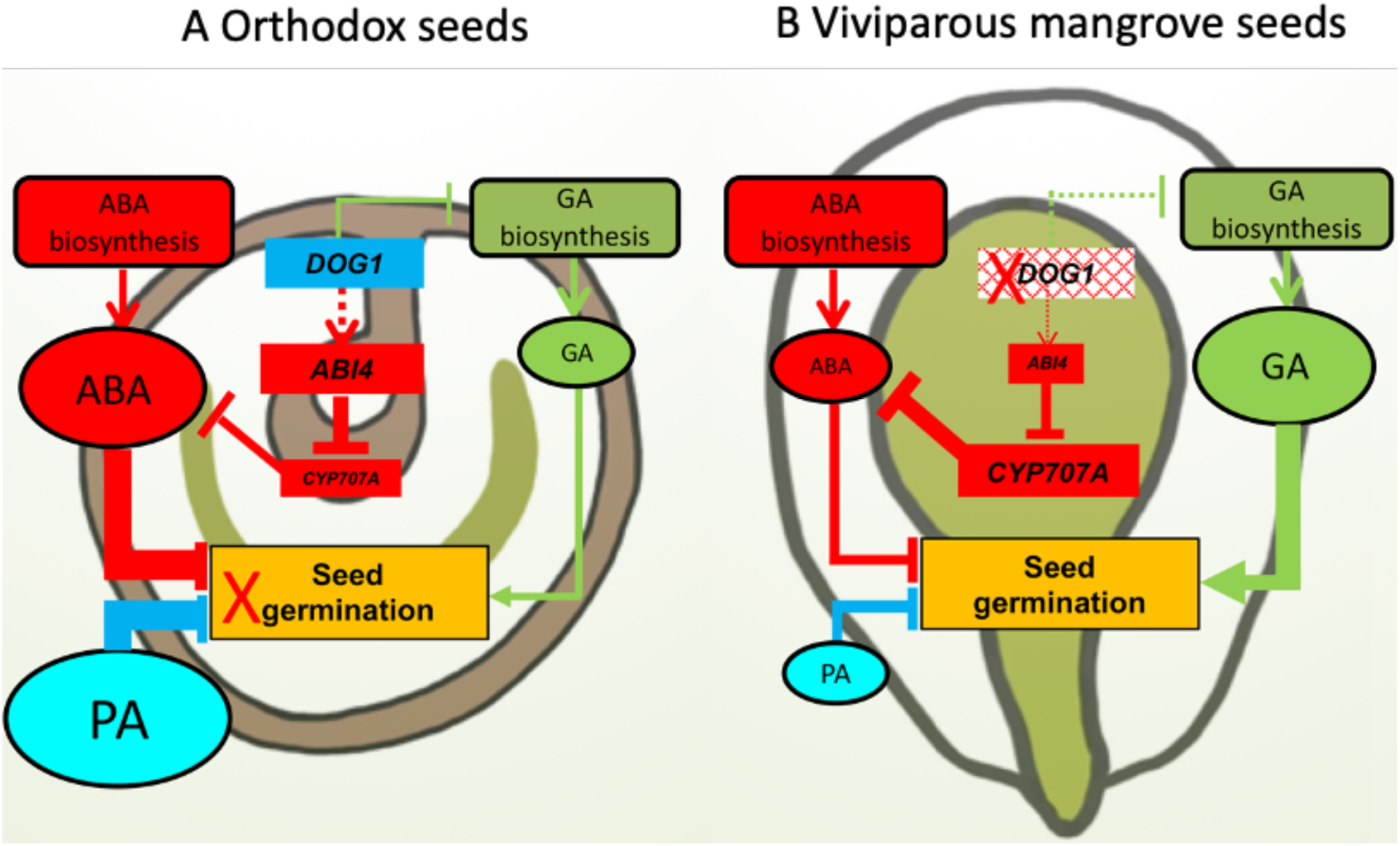
Schematic of non-vivipary and vivipary seed germination mechanisms in regard to *DOG1*. **A**, Regulation of seed germination in non-viviparous seed such as *C. brachiata*. **B**, Regulation of seed germination in the viviparous mangrove such as *K. obovata*. Red, green and blue represents ABA, GA and PA pathways, respectively. The sizes of texts and boxes represent relative level of gene expression between two models. Dotted arrow lines between *DOG1* and *ABI4* mean the observed but mechanically unclear regulation. Thickness of the lines represent relative strength of promotion or inhibition. In orthodox seed, *DOG1* promotes biosynthesis of ABA and PA but inhibits GA production, thus suppressing seed germination in the maternal plant. With *DOG1* is mutated in viviparous seeds, seed germination is promoted by more GA but less inhibition by ABA and PA.

## Methods

### Field sample collections

Individual plants of *K. obovata* in Zhangjiang Estuary National Mangrove Nature Reserve (Fujian, China; 23°53’45’’N-23°56’N, 117°24’7’’E-117°30’E) were chosen for genome sequencing. Fresh leaf, stem, flower, fruit and root tissues of *K. obovata* were harvested in the mornings from July to September 2015 and frozen immediately in liquid nitrogen. Genomic DNA was extracted from leaves using Fast Plant Genomic DNA Extraction System (#DP321, TIANGEN, China) and used to prepare sequencing libraries. Total RNAs of the dissected tissues were extracted using MiniBEST Plant RNA Extraction Kit (TaKaRa) and stored in a −80°C freezer for later use.

Leaves of *C. brachiata* were collected in June of 2016 from one healthy tree in the town of Gaoqiao, Zhanjiang City, Guangdong Province, China, and preserved in liquid nitrogen for DNA extraction. Tissues from three trees with similar age were collected for transcriptome sequencing from June 2016 to May 2017. Tissues of roots, stems, leaves, flowers, ovules, fruits, seeds, and embryos of these three trees were preserved in RNAlater (Ambion Inc., USA) for RNA extraction as mentioned above. Leaves and hypocotyls of *B. gymnorrhiza* and *C. tagal* were collected in August of 2019 in Wenchang, Hainan Province of China for DNA extraction.

### Genome sequencing, assembling and functional annotation

Detail method of genome sequencing, assembling and functional annotation are listed in the supplementary information. In brief, genomic DNA of *K. obovata* and *C. brachiata* are sequenced using PacBio, Illumina paired-end sequencing, and 10x Genomics sequencing platforms. These reads were then used for genome assembling mainly used using FALCON (https://github.com/PacificBiosciences/FALCON/) (Chin et al. 2016). For *B. gymnorhiza* and *C. tagal* genomes, Oxford Nanopore and Illumina paired-end reads were sequenced and then assembled using Wtdbg2 (Ruan and Li 2019). RNA-seq reads from various tissues of *K. obovata* and *C. brachiata* and homolog-based predictions were used for functional annotation of genomes.

### RNA and library preparation

For viviparous seed samples of *K. obovata*, MiniBEST Plant RNA Extraction Kit (TaKaRa) was used to isolate total RNA. The RNA-seq libraries were constructed following a refined SMART protocol (Hunt 2015). Briefly, mRNA was enriched with Oligo d(T)25 Magnetic Beads (New England Biolabs) and fragmented by heating at 94 °C for 3 min in 5× First Strand Buffer (Invitrogen). The fragmented mRNAs were reversely transcribed into cDNA by SMARTScribe reverse transcriptase (TaKaRa). After PCR amplification (Phire II), Illumina sequencing primer and sample specific barcode were added by ligation. About 300-500 bp DNA were selected using agarose gel electrophoresis and were purified by the Zymoclean gel DNA recovery kit (Zymo Research). High Sensitivity DNA Chips of Agilent Bioanalyzer 2100 were utilized to check library quality. Finally, qualified cDNA libraries were sequenced using the Illumina HiSeq 2500 platform at the core facility at College of the Environment and Ecology, Xiamen University. Tuxedo workflow using Hisat2-Stringtie-DESeq2 (Pertea et al. 2016; Love et al. 2014) was applied in quantification and normalization of RNA-seq gene expression.

### Data analysis

#### Quality control

Raw data (raw reads) of Fastq format were firstly processed through in-house perl scripts. In this step, clean data (clean reads) were obtained by removing adapters, reads containing ploy-N and low-quality reads from raw data. At the same time, Q20, Q30 and GC content of the clean data were calculated. All the downstream analyses were based on the clean data with high quality.

#### Reads mapping to the reference genome

Reference genome and gene model annotation files were downloaded from genome website directly. Index of the reference genome was built using Bowtie v2.2.3 (Langmead and Salzberg 2012) and paired-end clean reads were aligned to the reference genome using TopHat v2.0.12 (Kim et al. 2013), which can generate a database of splice junctions based on the gene model annotation file.

#### Quantification of gene expression level

HTSeq v0.6.1 (Anders et al. 2014) was used to count the read numbers mapped to each gene, and FPKM (Fragments Per Kilobase of transcript sequence per Millions base pairs sequenced) was used for normalization.

#### Differential expression analysis

Differential expression analysis of two conditions/groups was performed using the DESeq2 (Anders and Huber 2010). DESeq2 provide statistical routines for determining differential expression in digital gene expression data using a model based on the negative binomial distribution. The resulting *P*-values were adjusted using the Benjamini and Hochberg’s approach for controlling the false discovery rate. Genes with an adjusted *P*-value <0.05 found by DESeq were assigned as differentially expressed.

#### Tissue specific expression analysis

We divided 8 tissues into 2 categories, vegetative organs (roots, stems, leaves, flowers, and ovules) and reproductive organs (fruits, seeds, and embryos). Tissue specific expression was defined as that the expression level of organs was higher than the 80% of the total and indicated by τ index.

### RT-qPCR validation

For RT-qPCR, the First-strand cDNA was synthesized using a 5X All-In-One RT MasterMix (ABM, Canada). The relative quantification was done by ChamQTM Universal SYBR® qPCR Master Mix (Vazyme, China) from three biological replications, normalized to the reference actin gene from *K. obovata* and calculated by the 2-ΔΔCt method. Primer sequences are shown in Supplementary Table S6.

### Identification of missing genes for vivipary

We extensively collected 729 genes/gene families that are necessary for seed development, dormancy and germination (References were listed in Supplementary Table S4). Among them, 510 are *EMBRYO-DEFECTIVE (EMB)* genes which are essential for normal embryos development and were summarized previously (Meinke 2019). The rest includes genes related to metabolism, response and regulation of fundamental hormones, especially ABA and GA. Other genes reported to be associated with seed dormancy and germination were also included, such as transcription factors and chromatin modifiers.

The homologs of these genes in *K. obovata* and *C. brachiata* were searched by BLASTp with strict criteria of identity ≥ 40% and coverage ≥ 50%. For genes failed to pass these criteria, their reciprocal best hits were checked with E-value ≤ 1×10^-5^ and weighted coverage ≥ 0.33. Finally, the homologs of these genes/gene families in these two species were determined (Supplementary Table S4). In brief, three genes *RED1* (AT2G41945), *EMB2788* (AT4G27010), and *WRKY60* (AT2G25000) were found exclusively lost in *C. brachiata*. The function of the first two genes are unknown, and the function of *WRKY60* might be compensated by other *WRKY* genes. Four other genes, *EMB1789* (AT5G56930), *EMB3008* (AT5G39750), *AFP4/TMAC2* (*ABI FIVE BINDING PROTEIN 4*, AT3G02140), *DOG1* (AT5G45830) are commonly lost in *K. obovata and R. apiculata*. We searched *EMB1789, EMB3008, AFP4, DOG1* in other two viviparous mangroves (Supplementary Table S4), it showed that *EMB1789* had homologous genes as follow: ctg8 (*B. gymnorhiza*), ctg18 (*C. tagal*), and *EMB3008* was completely lost in four viviparous species. However, *EMBRYO-DEFECTIVE*(*EMB*) genes is a huge family, it’s difficult to decide the effects of *EMB1789* and *EMB3008* on vivipary. Although *AFP4* was not detected, its orthologous *AFP1, AFP2, AFP3* may be functional redundant of it. Therefore, *DOG1* is the only candidate that determines the seed reproductive pathway discrepancy between *K. obovata* and *C. brachiata*.

### Transgenic verification of *C. brachiata DOG1*

The genomic sequence of *C. brachiata DOG1* homologue (*CbDOG1*) was cloned into pCAMBIA1300S vector, and transformed to *Agrobacterium* then to *Arabidopsis dog1-5* mutant plants (SALK-022748) by floral dip method. *A. thaliana DOG1* (*AtDOG1)* was also cloned and transformed to *dog1-5* as a control. The transgenic T-2 seeds were used for the seed germination test. Freshly collected seeds (without cold treatment) were germinated on MS media in Petri dishes on top of Whatman paper, with 22°C/18°C temperature and 16 h/8 h light/dark cycles. Germination rates were taken after 10 days with 5 independent transgenic lines for each construct.

## Data availability statement

The datasets generated during the current study are available in the NCBI sequence archive. The accession numbers are PRJNA632974, PRJNA631086 and SRP261558.

## Acknowledgements

We thank Haidong Qu and Xiuxiu Wang for technical supports, students from Wenqing Wang’s lab for sample collection, and Taylor Li for language editing.

## Author contributions

Q.Q.L. and Y.S. conceived and designed the project. H.Q., X.Z., W.S. and S.He participated in library construction, sequencing and/or data analyses. P.J., C.Y. and X.Z. performed sequencing and bioinformatic analyses. W.H., M.F. and S.Hao performed RT-qPCR experiments. D.Y. performed transgene tests. W.W. collected and prepared samples. Y-Y.Z. involved in data analysis. Q.Q.L., Y.S., H.Q., X.Z., S.He, and C.Y. wrote the manuscript and all authors critically revised the manuscript.

## Supplementary Information

Additional Supporting Information may be found online in the Supporting Information tab for this article.

